# Geometry effects on protein mobility in a synapse

**DOI:** 10.1101/2025.01.12.632582

**Authors:** Simon Dannenberg, Sofiia Reshetniak, Sarah Mohammadinejad, Silvio O. Rizzoli, Stefan Klumpp

## Abstract

It is generally assumed that synaptic function requires a tight regulation of the mobility and localisation of synaptic proteins. Evidence for this hypothesis has been difficult to gather. Protein mobility can be measured via fluorescence recovery after photobleaching (FRAP), but the interpretation of the results remains challenging. In this study, we perform *in silico* FRAP experiments to study the effects of the synaptic geometry and/or protein binding to synaptic vesicles on protein mobility. We matched simulations with published FRAP data for 40 different synaptic proteins, to obtain diffusion coefficients, vesicle binding rates, and binding times. Importantly, we identify two mechanisms that govern the obtained recovery times: redistribution of material inside the synaptic bouton and inflow through the axon. We show that their dissection is crucial for the correct interpretation of FRAP experiments, especially for proteins binding to synaptic vesicles.

## I. INTRODUCTION

Signal transmission between neurons requires cellular connections termed synapses, where signals are mediated via the release and uptake of neurotransmitters, at the presynaptic and postsynaptic side, respectively. The neurotransmitters are stored in synaptic vesicles and their release requires complex cascade of chemical reactions, in which a plethora of different proteins are involved [1]. The major steps involve the fusion of the vesicles with the synaptic plasma membrane (exocytosis) and the subsequent retrieval of the involved proteins from the membrane (endocytosis) as well as their storage in the protein-rich synaptic vesicle cluster [2–5]. The abundance and the localization of the proteins involved in these processes is essential for synaptic function, and has been studied in much detail [6, 7]. Likewise, the functional roles of a large number of proteins has been studied extensively and is systematically annotated [1]. The picture that has emerged from these studied is that the synaptic vesicle cluster not only stores vesicles and provides a reserve of neurotransmitters, but also serves a hub for the proteins coordinating the synaptic organisation and dynamics on the pre- and postsynpatic side [8, 9]. In particular, the synaptic vesicle cluster gathers substantial amounts of soluble proteins [10], and thus provides a pool of these proteins, buffers their concentrations, and prevents their diffusive escape into the axon [10, 11].

The wealth of biochemical information about the proteins in the synapse is contrasted by rather scarce knowledge about the physical conditions in which these proteins perform their function. One aspect that is crucial here is the mobility of these proteins in the synapse as well as in the axon. The mobility of vesicles and other organelles in the synapse has been characterized [12–14], as has the mobility of some proteins [15–18]. Binding of synaptic proteins to synaptic vesicles has been characterized *in vitro*, for several synaptic proteins [19]. For a systematic assessment of protein mobility, Reshetniak et al. [20] catalogued the diffusion as well as their binding to synaptic vesicles for a set of 45 different proteins based on fluorescence recovery after photobleaching (FRAP) experiments [21]. The interpretation of the recovery times obtained from FRAP experiments is however not straightforward in complex geometries such as the synaptic bouton, where multiple factors can influence recovery times including binding of proteins to each other or to vesicles or organelles, the geometry of the underlying sample, and the localization of organelles (mitochondria, for example, may block the influx from the axon) [22, 23]. In addition, synapses are rather heterogeneous and can differ greatly in volume and shape as well as in the number of synaptic vesicles and the number and shape of large organelles [20]. The synaptic organization and the size of vesicle pools also changes as part of the synaptic plasticity [24]. This variability complicates the interpretation of FRAP data further, as typically recovery times are averaged over multiple experiments in different synapses.

Here we address these issues by simulating protein mobility in synapses of diverse shape and size. We perform *in silico* FRAP experiments to dissect different contributions to the recovery of fluorescence. First, we show that exact protocol of how photobleaching is conducted has a significant impact, in particular due to diffusion of the protein of interest during bleaching. We then identify two main processes which govern recovery of the signal: diffusive influx of proteins from the axon and redistribution of proteins inside the synapse. Comparing different synaptic geometries, we show that this effect can lead to fast-appearing recovery times, even in the absence of high mobility. Finally, we compare our simulations with the experimental data set of Reshetniak et al. [20], matching the simulation to the data to determine diffusion coefficients, binding times and binding rates for the proteins of their study. These comparisons show that, for most soluble proteins, fluorescence recovery is dominated by influx from the axon, albeit some proteins, as synapsin, differ from this behavior, through their strong vesicle-binding behavior.

## II. RESULTS

### A. Recovery times depend on photobleaching protocol in FRAP simulations

We first demonstrate the sensitivity of in silico FRAP experiments in synapses (or other confined cellular spaces) to the exact procedure used for photobleaching and data acquisition. To this end, we compared three different implementations of bleaching procedures that represent a spectrum of approaches to photobleaching in a confined geometry:

In the first two scenarios, bleaching occurs instantaneously either in the entire presynaptic region or only in a small (400nm diameter) subsection of it. In the third case we followed an experimental protocol of Reshetniak et al. [20]. We bleached again only a small subsection but did so continuously for 80 ms. This allows proteins to diffuse in and out during the bleaching effectively enlarging the bleached region.

In all three scenarios, we considered the same synaptic environment and modelled the mo- bility of a freely diffusion protein with a constant diffusion coefficient according to a reaction diffusion master equation (for details see Methods). However, the three scenarios exhibit substantially different recovery behavior as shown in Figure 1A by series of simulated fluo- rescence images for the three scenarios (note the different time scales) and in Figure 1B by the corresponding recovery curves.

**FIG 1.**
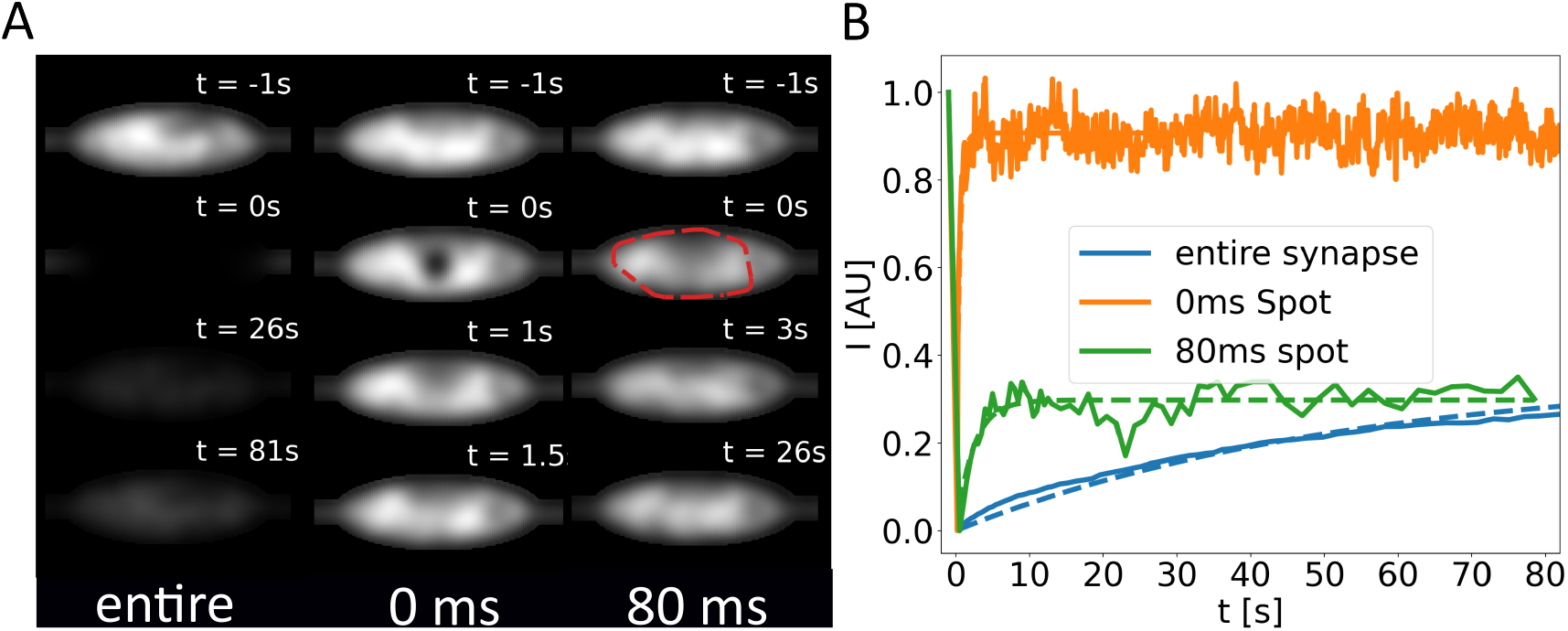
**A** Synthetic microscopy images from simulated FRAP experiments in the presynaptic region with three different photobleaching protocols. In the first protocol the entire synapse is photobleached instantaneously, while in the second and third only a portion is photobleached, either instantaneously or over a time interval of 80 ms. Note the different time stamps. **B** Recovery of fluorescence intensity in the region of interest of the different protocols. For the first cases the regions of interest are the spots where the photobleaching was applied, in the third case, the region of interest, marked by the red circle, is determined afterwards according to a threshold criterion (see Methods).

In the first scenario, where the entire presynaptic region is bleached instantaneously, all proteins within the simulated presynaptic region are labelled as non-fluorescent at the time of bleaching. Afterwards, we selected the entire synapse as our region of interest (ROI) to record the recovery. This was done until the equilibrium concentration was reached again. Subsequently, the recovery curve was fitted to a single exponential *I*(*t*) = *I*_0_(1 − *e*^−τ*/t*^), from which the recovery time τ can be extracted. The resulting recovery curve from the simulation and the corresponding fit are depicted in blue in Figure 1B.

In the second scenario, where only a spot within the synapse is bleached instantaneously, we designated a cylindrical section of the synapse (at the synapse’s center and with a 400 nm diameter) as the bleached volume, mimicking the path of the bleaching laser through the synapse. The size and shape of the bleached region was chosen to correspond to bleaching in experiments with fixed proteins in a synapse [20]. Subsequently, we used the same region also as our ROI in which we monitor the recovery of fluorescence and determined the recovery curve, depicted as the orange curve in Figure 1B.

For the third protocol, bleaching a spot over 80 ms, we chose to mimic the procedure used in the study by Reshetniak et al. [20]. Most importantly, we took into account that bleaching is not instantaneous, but that the bleaching laser is applied for 80 ms on a spot at the synapse center (described in the same way as in the second protocol). During the bleaching process, proteins diffuse in and out of the laser spot, leading to an increase in the effective bleaching spot size. Crucially, this enlargement of the spot depends on the diffusion coefficient of the proteins, as faster diffusion leads to a wider spread of the spot. To identify the region in which the recovery should be monitored, we followed the protocol used in the experiments [20] and selected all pixels with a significant change in intensity (marked by the red circle in Figure 1A) as the bleached region and as the ROI for monitoring the recovery (see Methods). The resulting recovery curve is shown in green in Figure 1B. For all three cases, the recovery curves shown were normalized to their intensity values prior to bleaching. This is important as in the three scenarios significantly different total amounts of protein are bleached, as evident from the synthetic images.

Comparing the three resulting curves, we can see clear differences in their qualitative shape which translate into a drastic differences in recovery times (52 s vs. 0.5 s vs. 2.3 s, respectively). Moreover, recovery is almost complete in the case of instantaneous bleaching of a spot, while it is only partial in the other two cases (within the 80 s of simulated recovery). These differences can be interpreted as follows: If the entire synapse is bleached, the recovery curve displays a slow, continuous increase in intensity that is still ongoing at the end of the simulated time period. This is mediated by diffusive transport of proteins from the axons into the synapse, as there are no unbleached proteins left in the synapse. By contrast, when only a spot in the synapse is bleached, recovery to the full intensity from before photobleaching is very rapid, as it occurs through redistribution of proteins inside the synaptic volume. Strikingly, the recovery curve for continuous bleaching for 80ms falls in between the previous two. An initial sharp increase in intensity is accompanied by an (apparent) plateau over the remainder of the simulation. As only small part of the synapse is effectively bleached, recovery can again occur through local redistribution. However, a relatively large immobile fraction can be seen in this case, as influx of unbleached material happens at a slower pace. This is due to the fact that significantly more material is bleached compared to the case of instantaneous photobleaching, although still much less than in the scenario where the entire synapse has been bleached.

These observations demonstrate that the detailed protocol of how photobleaching is simulated (or, likewise, carried out experimentally) can significantly impact the results by changing the dominant mechanism of recovery, modulating between redistribution of material inside the confined space of the synapse and inflow into that space from the axon.

### B. Heterogeneity of synaptic geometries results in heterogeneous recovery times

Our analysis so far shows that the choice of (in silico) protocol that is used to implement FRAP affect the measured recovery times. On top of that the geometry of many biological systems is very heterogeneous. A prime example are synapses. For example, their volume and the number of synaptic vesicles they contain can vary by an order of magnitude, as shown in Figure 2 based on data from ref. [20]. Moreover, their shapes are known to undergo significant changes upon stimulation. Thus, in addition to the protocol how the FRAP signal is recorded, also the underlying geometry is expected to affect the recovery times. To investigate this effect, we generated an ensemble of 30 different presynaptic regions. These are modeled as ellipsoids of different sizes, aspect ratios, containing different numbers of mitochondria, vacuoles, and vesicles (for details, see Methods). The histograms in Figure 2 show examples of the data used to generate the synapse models.

**FIG 2.**
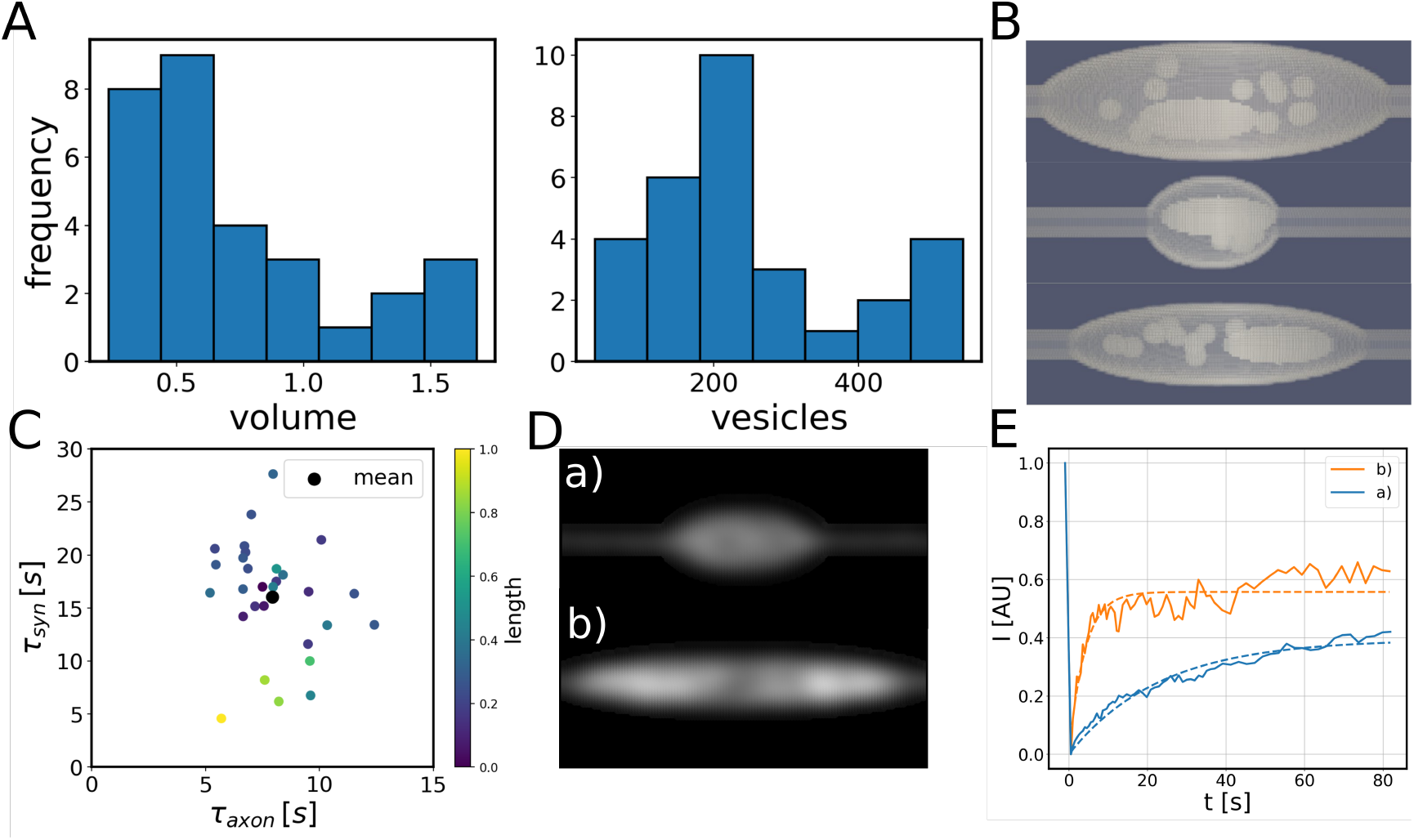
**A** Histograms of synaptic geometry characteristics used to create in silico synapses (data from ref. [20]). **B** Examples of synthetic geometries. The visible structures within the ellipsoids represent excluded volume occupied by mitochondria and vacuoles. **C** Variability of recovery times after photobleaching in the axonal and synaptic regions. Each dot represents a different synaptic geometry. Simulations were done for pure diffusion with D = 0.1 *µm*^2^*s*^−1^ and no binding to synaptic vesicles. **D** Synthetic microscopy images taken 0.5s after photobleaching for two different synapse geometries and **E** corresponding recovery curves.

To analyze the influence of the synaptic geometry, we first studied again the case of purely diffusing proteins and simulated FRAP experiments in the different geometries from the ensemble of synapse models, keeping the a fixed diffusion coefficient fixed at D = 0.1 *µm*^2^*s*^−1^. For all geometries, we conducted FRAP in the synapse and in the axon. The geometry of the axon is the same in all models. Thus, for every synapse geometry, we obtained a pair of recovery times (τ_*syn*_, τ_*axon*_).

These resulting recovery times are plotted in Figure 2, where every blue point shows one synapse geometry. The orange point indicates the mean after averaging over all experiments. Note that we used different scales on the axes, as recovery is considerably slower in the synapse. Importantly, our results show a wide range in recovery times in the synapse, varying between 5s and 30s, while the recovery times in the axon values are more uniform. This can be explained by comparing the two individual synapses highlighted as *a* and *b* in Figure 2 that show rapid and slow recovery, respectively. In Figure 2D we show simulated fluorescence images that display the intensity of the first image of the sample taken after bleaching, at *t*_*b*_ = 0.5 s. These two presynaptic regions are quite different in volume and length. Synapse *a* is significantly more elongated, so only its central region is bleached, while synapse *b* is bleached entirely. This difference resembles the difference between bleaching protocols studied above: For synapse *b*, the only path for fluorescence recovery is influx of fluorescent protein from the axon. This results in an approximately single-exponential recovery curve as shown in Figure 2E. In synapse *a* however, material can be redistributed within the synapse, which provides a second, faster recovery mechanism. As a consequence, we observe a jump in the fluorescence recovery signal in the beginning of the recording, accompanied by a slower rise at later times. Therefore, the combination of the two diffusive mechanism observed above leads to a fluorescence recovery curve with two timescales. Importantly, this difference between the two recovery curves is exclusively due to differences in the synaptic geometry, as the molecular motion is the same and purely diffusive in both cases.

### C. Recovery times reflect a balance of redistribution within the synapse and inflow from the axon

Our analysis so far indicated two different modes for fluorescence recovery in the synapse: influx from the axon and redistribution inside the presynaptic region. To assess this picture more quantitatively and to test how generally applicable it is, we selected different diffusion coefficients between 0.1 and 20 *µ*^2^*s*^−1^ and for each simulated FRAP experiments in 30 synaptic geometries. To compare the results between different synapses, we rescaled the synapse length *L* with the distance the protein of interest diffuses during bleaching, 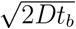. The effective length 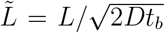characterizes how much of the synapse is bleached. If the diffusive distance is greater than the synapse length, 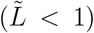, we expect the entire synapse to be bleached. On the other hand, if 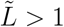only a part of the synapse is bleached. Therefore, this effective length allows us to qualitatively assess which of the two mechanisms of fluorescence recovery should be dominant.

In Figure 3, we plotted the recovery times in the synapse from all simulations (different geometries and different diffusion coefficients) against the effective synapse length 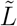. The diffusion coefficients used in the simulations are indicated by the color of the data points. Strikingly, the data points for different synapse geometries and different diffusion coefficients fall approximately on one curve. This curve displays a prominent maximum for 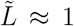. This maximum separates two regimes, in which the recovery time increase and decreases, respectively, as a function of the rescaled synapse length 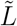. Since the dependence on the diffusion coefficient is included in the rescaled synapse length 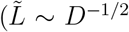, this also means that the recovery time is not monotonic as a function of the diffusion coefficient. For rapid diffusion 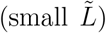, the conventional picture where shorter recovery times correspond to higher mobility is valid. However, this is not the case for intermediate to slow diffusion 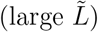, where only a fraction of the synapse is bleached and redistribution in the synapse contributes to the recovery process. Varying the diffusion coefficient in this regime not only changes the mobility of the protein, but also the relative weights how much the two recovery processes contribute to the observed recovery dynamics. For very slow diffusion 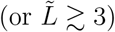, we expect a third regime, where recovery occurs exclusively by redistribution in the synapse.

**FIG 3.**
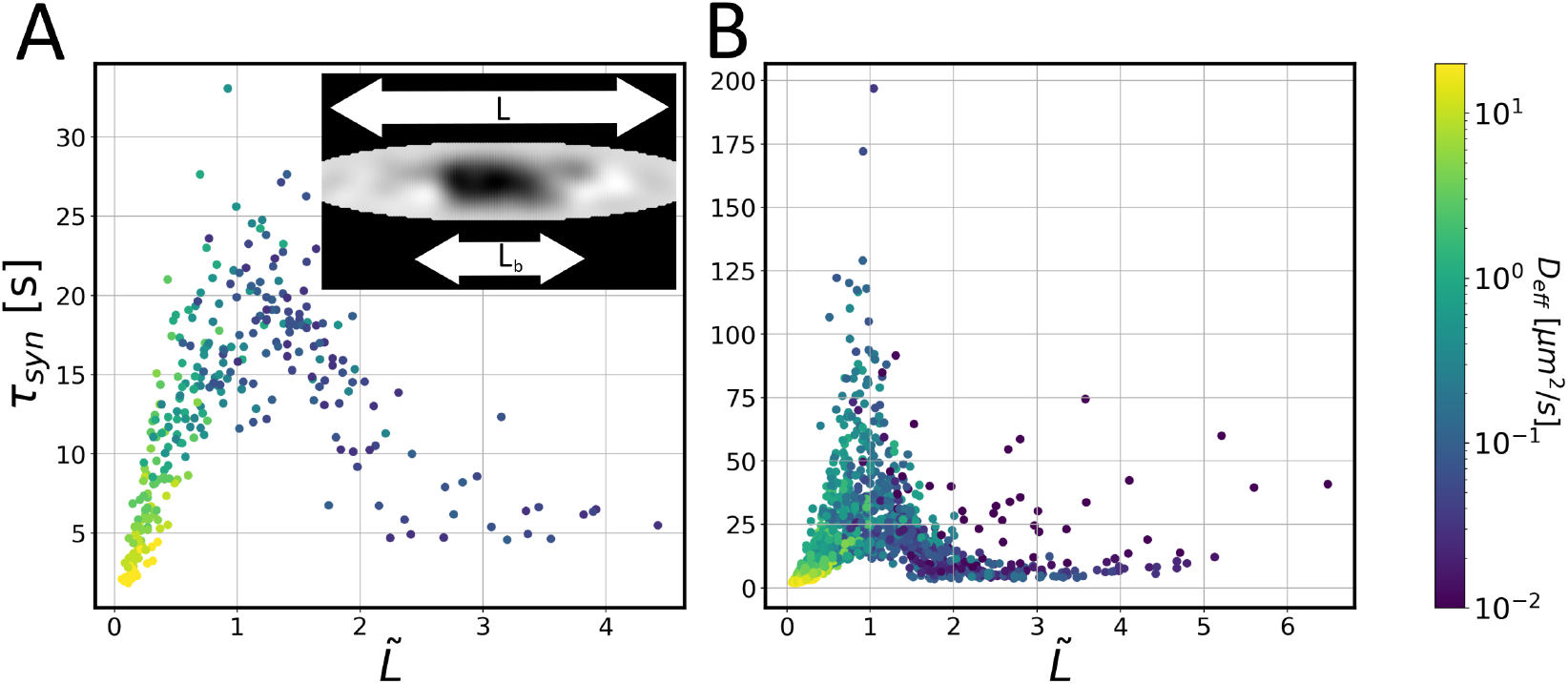
**A** Recovery times in different presynaptic geometries plotted over the effective synapse length (synapse length normalized to the distance traveled by a protein during photobleaching). The color code indicates the diffusion coefficient. **B** Same as in A but including binding to vesicles and unbinding and introducing an effective diffusion coefficient to determine the effective synapse length. Insets show the same synaptic geometry after bleaching but using different binding rates.

In this regime, τ_*syn*_ should increase again with increasing 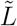, but this regime is not explored in our data.

These observations underline that the recovery time is governed by a combination of synapse geometry and the diffusion coefficient of the protein. But what happens when proteins do not diffuse freely but have more complex dynamics? To answer this question, we included binding to synaptic vesicles into our model and simulated FRAP in different synaptic geometries with a range of diffusion coefficients, binding rates and unbinding rates. Binding transiently renders proteins immobile and thus reduces their mobility. To apply the same analysis as above, we replaced the diffusion coefficient of the proteins with an effective diffusion coefficient and modified the rescaled synapse length 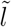 to depend on that effective diffusion coefficient, 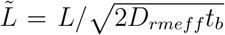. The effect of binding to vesicles is incorporated into the effective diffusion coefficient *D*_eff_ as a reduction of mobility by a factor given by the probability that the protein is not bound and diffuses freely. This leads to 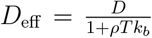(see **??** for details). Here, *ρ* is the vesicle density, *T* is the average time a protein is bound to the vesicle and *k*_*b*_ is its binding rate. Importantly, this approximation requires that binding and unbinding occur on timescales faster than the time of the experiment. With this generalization of the rescaled synapse length we plotted the recovery times in the synapse from simulations with binding/unbinding as function of this modified 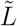, as shown in Figure 3B. Importantly, we observe the same qualitative behavior as for pure diffusion with the same two regimes. The collapse of the data points into one curve is not perfect and the data scatter around that curve, in particular when binding/unbinding is slow and the mean field approximation of an effective diffusion coefficient is less accurate. Nevertheless, the qualitative picture is the same as without binding/unbinding. As before, for longer synapses with 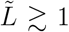, the redistribution of material within the synapse becomes important and the recovery time is reduced with increasing 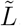.

### D. Qualitative Roadmap to access protein mobility from FRAP experiments

Our previous analysis has shown that a short recovery time is not necessarily the result of high mobility, as different recovery times may reflect different recovery mechanisms rather than different diffusion coefficients. This effect is illustrated in Fig. 4A. Here we highlighted two sets of simulations done in many different synaptic geometries. One for a freely diffusing protein with diffusion coefficient *D* = 0.5*µm*^2^*s*^−1^ (orange) and one for a protein that diffuses with the same diffusion coefficient, but also binds to vesicles (*D* = 0.5 *µm*^2^*s*^−1^; *T* = 1*s*; *k* = 1000*s*^−1^, black) and thus has a smaller effective diffusion coefficient of *D*_eff_ = 0.09 *µm*^2^*s*^−1^ (note that unbinding and unbinding rates are sufficiently high, so the description by an effective diffusion coefficient is accurate). Averaging over the different synaptic geometries results in recovery times of τ_*syn*_ = 16.2 *s* and τ_*syn*_ = 13.2 *s* for the freely diffusing and the vesicle-binding protein, respectively. Thus, a smaller recovery time is found for the latter protein even though its effective diffusion is slower. This somewhat paradoxical observation can be understood by realizing that the two cases fall on different branches of the recovery time master curve (Fig. 4A) and that their time scales cannot be directly compared as they reflect different recovery mechanisms, influx from the axon for the freely diffusing protein and redistribution in the synapse for the vesicle-binding protein.

**FIG 4.**
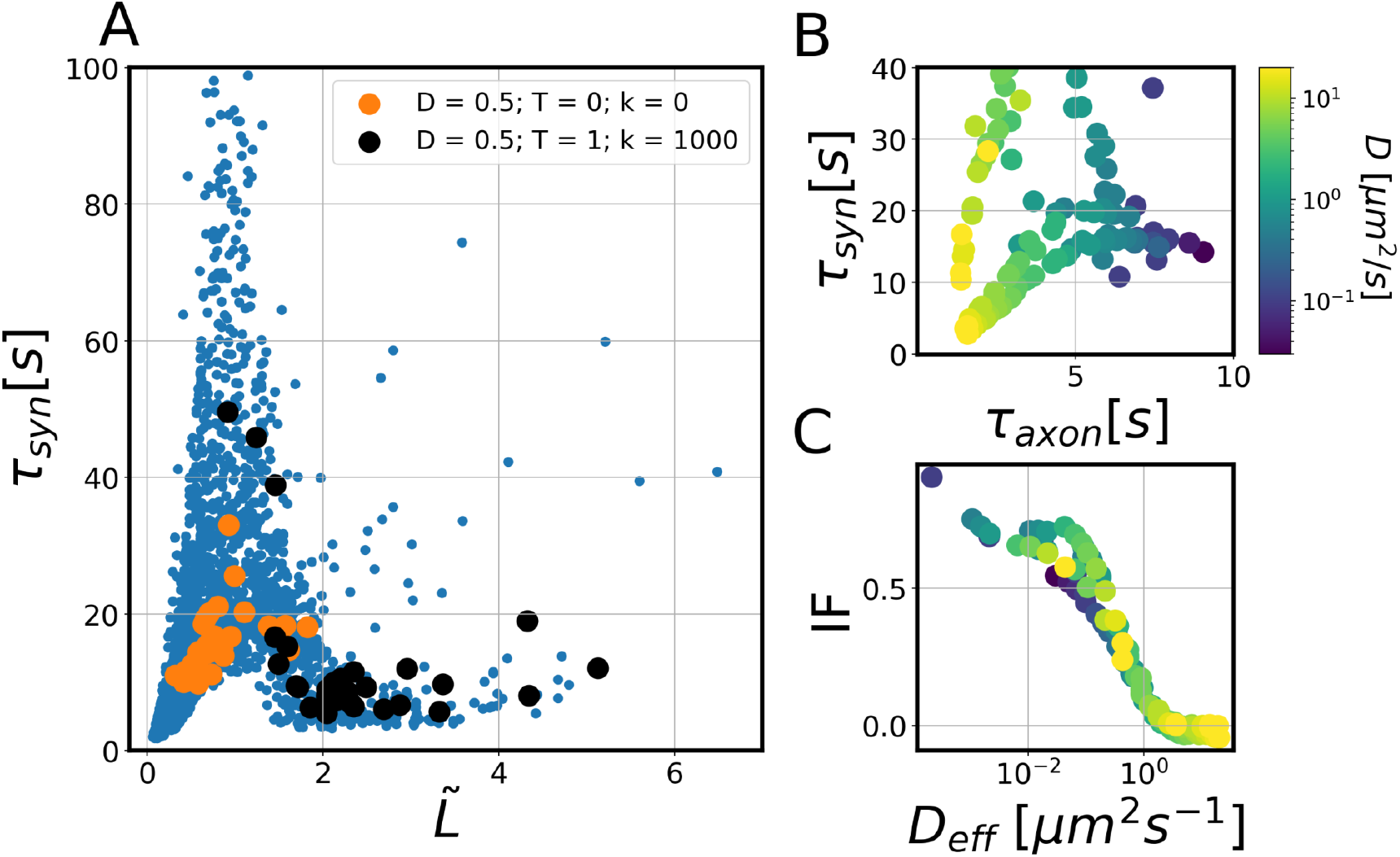
**A** Recovery time in the synapse τ_*syn*_ vs. the effective synapse length (synapse length normalized to the distance a protein diffuses during photobleaching) for different parameter com- binations and geometries. Highlighted in orange and in black are two specific parameter sets representing two hypothetical proteins with different properties (*D* = 0.5*µm*^2^*s*^−1^, orange, and *D* = 0.5*µm*^2^*s*^−1^; *T* = 1*s*; *k* = 1000*s*^−1^, black). **B** Scatter plot of recovery times in synapse and axon. The diffusion coefficient used in the simulation is color-coded. C) Immobile fraction plotted against the effective diffusion coefficient. Here, we binned the data according to the effective dif- fusion coefficients and averaged the immobile fraction.

To distinguish the two cases, additional information is required. One such addition input is given by the recovery time in the axon, which as a thin quasi-one-dimensional compartment is much less affected by the geometric heterogeneity. Indeed, the color coding by the underlying diffusion coefficient in Fig. 4B shows a pronounced correlation between the diffusion coefficient and the recovery time in the axon. A second additional input is the immobile fraction measured in the synapse, which in contrast to the recovery time shows a monotonous dependence on the (effective) diffusion coefficient. In the following, we will include both observable as input for determining the dynamic parameters of specific proteins from FRAP experiments.

### E. Fitting diffusion coefficients, binding times and binding rates to experimental data

Based on our previous analysis that established the need for at least two observables to determine the molecular parameters from FRAP recovery curves, we finally used our simulations to fit the experimental data from ref. [20]. To determine diffusion coefficients and binding and unbinding rates for the set of proteins studied in that work, we included the recovery times τ_*axon*_ and τ_*syn*_ and the corresponding immobile fractions IF_syn_, IF_axon_ into our fit (see Methods). An overview of the pairs of experimental recovery time for all studied proteins is shown in Fig. 5A, (for the corresponding immobile fractions, see Fig. 7). Three proteins in the data set were only detected in the synapse and not in the axon, these were omitted here.

**FIG 5.**
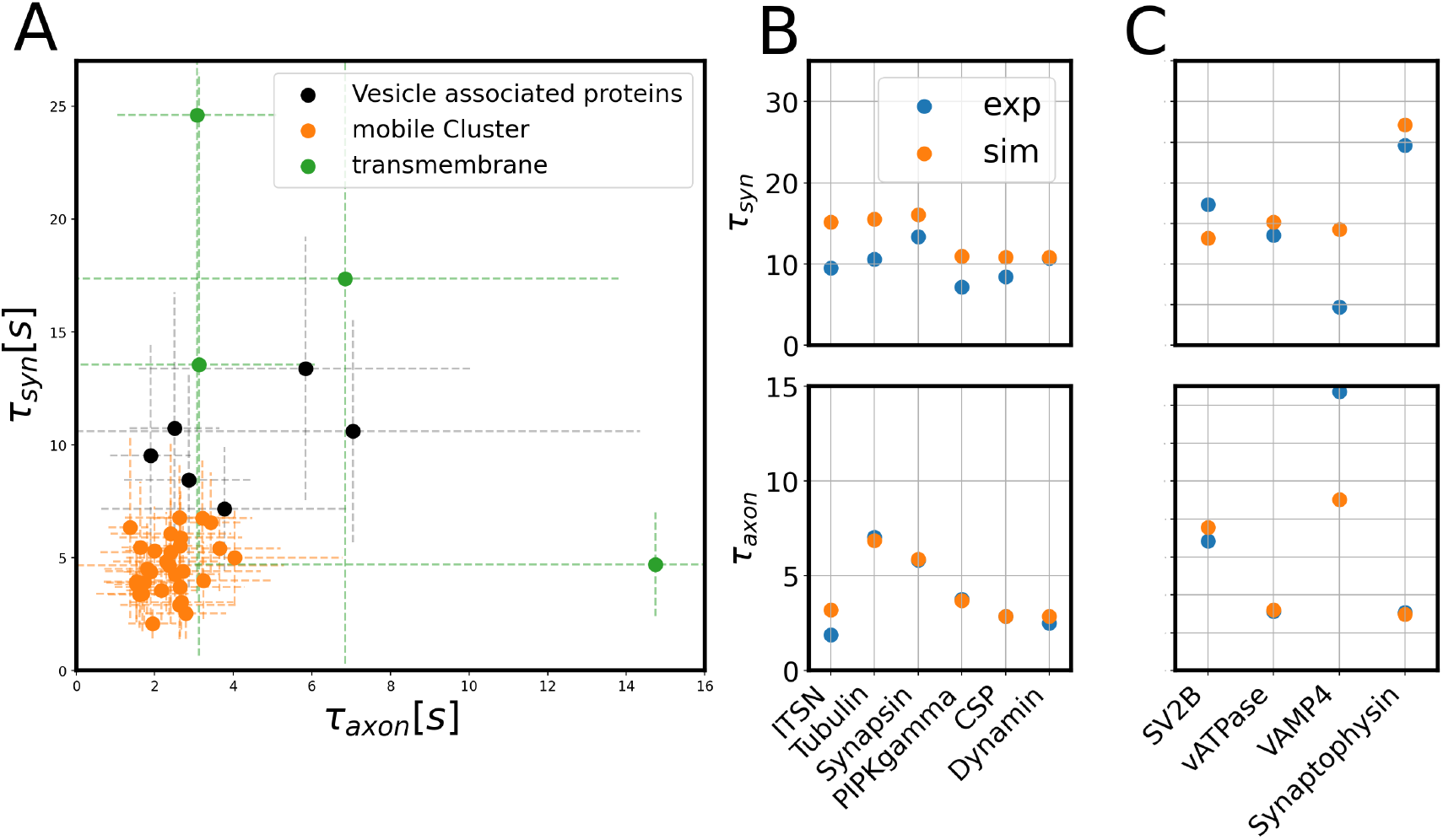
**A** Pairs of experimentally recorded recovery times in the axon and the synapse (τ_axon_, τ_syn_) for 42 proteins (data from ref. [20]. The proteins shown in orange form a cluster of highly mobile proteins (identified using a K-means and a Gaussian Mixture Model) for which mobility is not detectably reduced by interactions with synaptic vesicles. **B**,**C** Best fitted results matching our simulated recovery times in the times in the axon and in the synapse (orange) to the corresponding experimental results (blue). B shows soluble and vesicle-associated proteins, C transmembrane- proteins.

**FIG 6.**
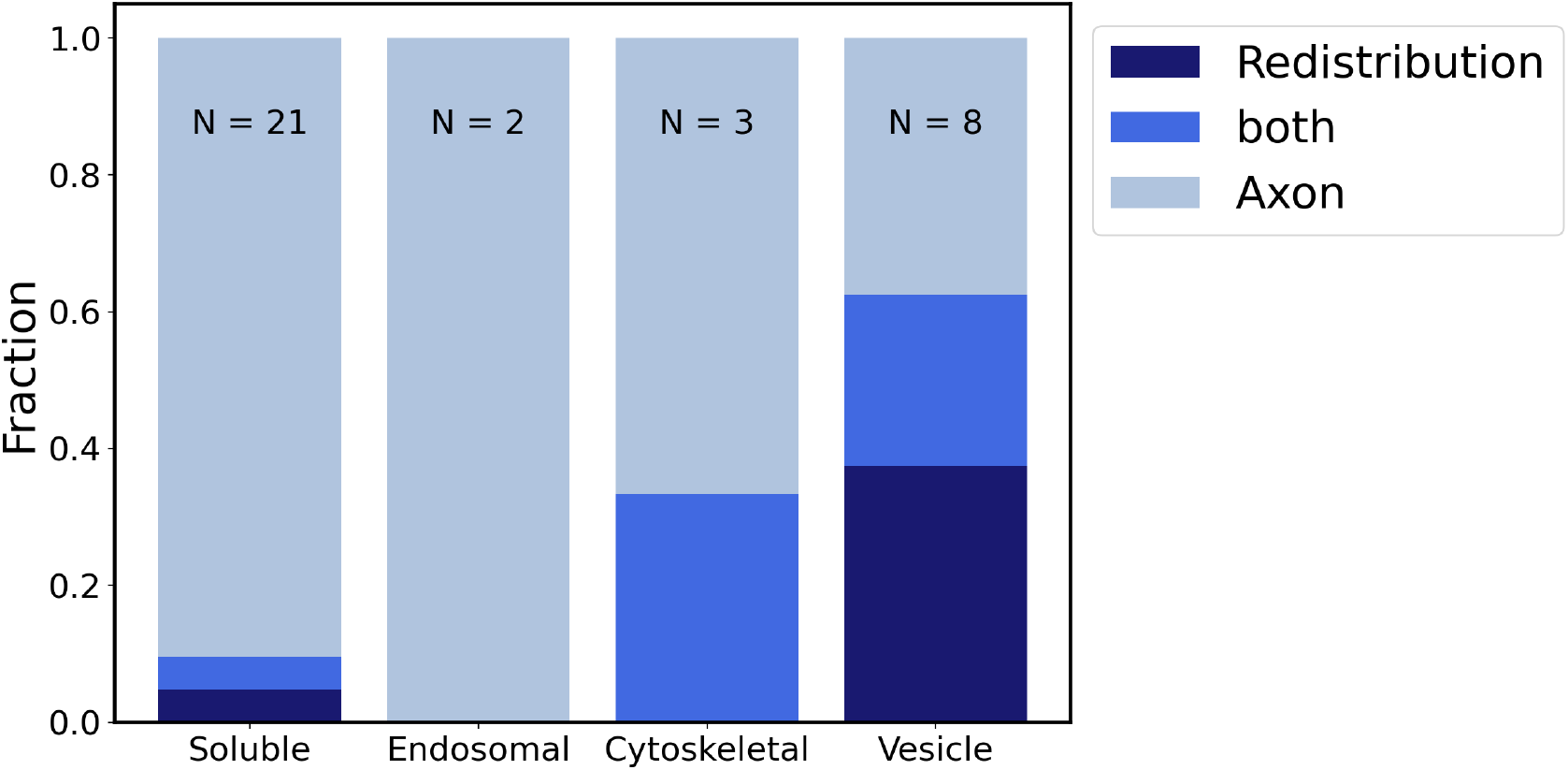
Statistics of the dominant fluorescence recovery mechanisms (redistribution in the synapse vs. influx from the axon) in different protein categories: Fractions of proteins using either mechanism. For proteins denoted as ‘both’, both mechanisms contribute to a similar degree.

The large variability between measurements in different synapse, indicated by the standard deviations of the recovery times, makes it difficult to distinguish the kinetics of the proteins with short recovery times from a purely diffusing protein that does not interact with synaptic vesicles. Therefore, we used a K-means algorithm and a Gaussian Mixture Model to separate between proteins that can be distinguished in terms of their mobility properties. Both algorithms identified a “rapid recovery” cluster of 30 proteins whose recovery times are centered around τ_*axon*_ ≈ 3*s* and τ_*syn*_ ≈ 5*s* (orange points in Fig. 5A, the identified cluster is independent of the number of centers set in the clustering procedure). Most of these molecules are soluble in the synaptic cytosol, although some are membrane-bound (membrane-anchored EGFP, SNAP23, SNAP25, Syntaxin 1), while two are bona fide synaptic vesicle proteins, and therefore should be found mostly within the slow-moving vesicles (VAMP2, Synaptotagmin 7). While our model predicts that such proteins have limited interactions with the vesicle cluster (in the sense that these interactions do not significantly slow down their diffusion), this is obviously not the case for VAMP2, present at ∼ 70 copies per synaptic vesicle. Other molecules in this cluster are also known to bind strongly to synaptic vesicles, including alpha-synuclein [25], or Rab3 and Rab5 [26]. Many other proteins in this cluster are bound to and buffered by vesicles [10]. Nonetheless, our model cannot determine their vesicle-interaction parameters (as their diffusion appears unaffected by the interaction). In principle, their diffusion coefficients can still be estimated from their recovery time in the axon. The resulting values are listed in Supporting Table S1, albeit the behavior of these molecules in the synapse remains unclear.

The remaining 10 proteins, whose behaviors differ from those in the rapid-recovery cluster, contain 6 soluble or vesicle-associate proteins (shown in black in Fig. 5A) and 4 transmembrane proteins (shown in green). For the soluble and vesicle-associated proteins, we determined all parameters (diffusion coefficient, binding and unbinding rates) from the recovery times and immobile fractions in the axon and the synapse. The recovery times resulting from the best fits are shown in Fig. 5B and the corresponding immobile fractions in supporting figure 8. The resulting values of all parameters are listed in Table I; they are also included in Supporting Table S1.

**TABLE 1.** Overview of the results of fitting FRAP data for individual proteins with the simulation.

| Protein | category | $D$<br>$\mu\text{m}^2\text{s}^{-1}$ | $k$<br>$\text{s}^{-1}$ | $T$<br>$\text{s}$ | $D_{\text{eff}}$<br>$\mu\text{m}^2\text{s}^{-1}$ | Recovery<br>mechanism |
| --- | --- | --- | --- | --- | --- | --- |
| Intersectin 1-L | soluble | 1.00 | 1000.0 | 1.0 | 0.021476 | Redistribution |
| $\alpha$ Tubulin | cytoskeletal | 0.30 | 0.0 | 0.0 | 0.300000 | Both |
| Synapsin 1A | soluble | 0.70 | 10.0 | 0.1 | 0.669495 | Both |
| PIP5KIγ | soluble | 3.00 | 10.0 | 0.1 | 2.869263 | Axon |
| Dynamin | soluble | 5.00 | 100.0 | 0.1 | 3.434900 | Axon |

Additionally, fitting the data for the 4 transmembrane proteins, our simulations predicted a diffusion coefficient comparable to the diffusion coefficient of synaptic vesicles for three of these proteins (SV2B, vATPase, synaptophysin; the resulting fits shown in Fig. 5C). Together with the high immobile fraction measured in these experiments, this led us to believe that their recovery measurement was mostly influenced by vesicle diffusion. As a negative control, we also considered a fit for VAMP4, the fourth transmembrane protein in the dataset, whis is present in endosomes that are trafficked along axons, to and from synapses [27] implying that our model should not be applicable. Indeed, the quality of the fit is considerably poorer than for the other proteins.

The recovery times obtained by from the best-fitting simulation match the experimental recovery times very closely for nearly all proteins.

The immobile fraction in the axon is systematically higher in the experiments than in the simulations with a ‘noise’ floor of around 20 percent that is not necessarily associated with vesicle binding or slow mobility as considered in our simulations. The excess immobile fractions on top of that ‘noise’ floor matches our simulations rather well. It is also worth mentioning that fitting the immobile fraction in the synapse is reminiscent of a binary decision, where our fit basically distinguishes between binding and non-binding proteins.

The resulting values of the diffusion coefficients are consistent with earlier work. The diffusion coefficient of synaptophysin has been reported to be 0.049 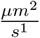[15] in excellent agreement with our result of 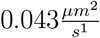. The diffusion coefficient for synapsin in the axon has been reported to be 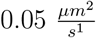and to be 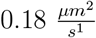inside the synaptic bouton, slightly lower than our obtained values of 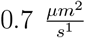 and 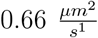, respectively. For mGFP, values of 15 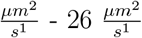[28] have been reported. However, also much slower diffusion was obtained in measurements in the dendritic shaft resulting in a diffusion coefficient 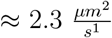[29]. Our fit suggests 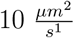, well within the range.

For proteins outside the “rapid recovery” cluster, we obtain large binding rates and binding times (Table I), reflecting strong association with vesicles. These parameter combinations result in small effective diffusion coefficients, e.g. *D*_eff_ = 0.0.02*µ*m^2^*/*s for Intersection 1-L. This value is comparable to the diffusion of the vesicles themselves (0.01*µ*ms^−1^ REFS, which is not explicitly considered here, but will contribute to the redistribution of the proteins.

Finally, we come back to the two recovery mechanisms identified above: For each protein, Table I also lists the dominant recovery mechanism. For proteins in the “rapid-recovery” cluster, we suggest that recovery by influx from the axon (Table S1) is an important mechanism. An overview of the recovery mechanisms is shown In Fig. 6, where we plot the fractions of proteins in the different protein categories that exhibit predominantly recovery by influx from the axon, by redistribution in the synapse or a combination of both. For most soluble proteins, recovery happens via influx from the axon. A notable exception is synapsin, which is strongly associated with vesicles as reflected in a large binding rate (Table I) and shows fluorescence recovery by redistribution as well as from the axon.

## III. CONCLUDING REMARKS

Our simulations demonstrate the sensitivity of modelling FRAP experiments on the protocol used for photobleaching. Furthermore, using different geometries we show, that in the case of presynaptic boutons and similar geometries a two timescale behaviour of the recovery curve does not mean a non-diffusive dynamic, but rather two distinct mechanisms for fluorescence recovery, namely redistribution of protein within the synapse and influx from the axon. The prevalence of these two mechanisms is different in soluble and vesicle- or endosome-associated proteins. For the balance of these two mechanisms, the fraction of the synapse that is photobleached is crucial and that fraction in turn depends, for example, on the size of the synapse, but also on the extent to which the fluorescent proteins diffuse during photobleaching. We observed that the geometry effects on FRAP can be well de- scribed as dependent on the ratio of the synapse length and the distance traveled during photobleaching. These results are likely also to apply to other systems with similar geometries, specifically to postsynaptic proteins where redistribution within dendritic spines may similarly compete with influx from the dendrite.

In practice, strong geometry effects can be avoided by selecting large synapses for FRAP experiments or by reduzing the area that is photobleached. The effective synapse length provides a criterion 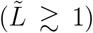how important geometry effects will be. If they cannot be avoided, interpretation of FRAP results can be based on a combined analysis of recovery time, bleached area, and immobile fraction to distinguish between the different regimes or recovery mechanisms in the geometry effect.

Overall, our analysis does not explain the behavior of several fast-recovering molecules, including several soluble proteins, but also major components of synaptic vesicles (VAMP2, Rab3), of the largely immobile synaptic cytoskeleton (actin), or of the plasma membrane (syntaxin 1, SNAP25). Further analyses will be needed to explain their behaviors in the future.

## IV. METHODS

### A. In silico representations of presynaptic regions

We approximated synapses as ellipsoidal structures which have two openings at the end of the long axis at which they are connected to the axon. The axon is modeled as a cylindrical tube of 200nm diameter and 5*µ*m length [30] which is approximately the mean value for neurons cultured from mice as used in the experiments by Reshetniak et al. [20]. The remaining parameters governing the geometry of the synapse are: length and aspect ratio of the synapse, number of synaptic vesicles, numbers of vacuoles and mitochondria and volume occupied by all of them. These parameters are extracted from EM images of 30 synapses creating a representative distribution of synaptic geometries [20]. The geometries are represented in our simulations with a spatial resolution of 25 nm, which is equal to the one of the EM measurements on which the synaptic geometries are based. We modelled mitochondria and vacuoles as occupied space inaccessible to the diffusing proteins; vesicles are represented as specific regions that allow for binding. They are placed randomly inside the bouton.

### B. Simulations of protein mobility

We simulate protein mobility in the presynaptic geometry using the next subvolume method [31, 32], a spatial variant of the Gillespie algorithm for stochastic simulations [33] that solves a reaction-diffusion master equation (RDME) on a discrete lattice. Diffusion is represented as a reaction that describes transfer of proteins (hopping) between neighboring voxels with a rate that is dependent on the diffusion coefficient *D* (hopping rate given by 6*D/a*^2^ with the voxel size *a*). In addition to diffusion, our model accounts for the binding and unbinding of proteins to and from vesicles. The time points at which hopping, binding and unbinding occur are determined via a Gillespie algorithm. Specifically, the next subvolume method combines two variants of that algorithm, the ‘next reaction method’ [34] to determine the subvolume in which the next reaction will occur and the ‘direct method’ [33] to determine that reaction within that subvolume. With this approach we monitor the protein concentration profile over time.

The synaptic geometries are incorporated into the lattice as a confinement of the proteins such that hopping is only allowed within the accessible geometry. In addition, the geometry designates certain voxels as containing vesicles, where binding and unbinding occur with rates *k*_*b*_ and 1*/T* (with the retention time *T*, the time spent on average bound to a vesicle), respectively.

We used periodic boundary conditions at the two end of the axon, such that proteins reaching the end on one side of the axon reenter on the other side. This mimics the exchange with a reservoir of proteins provided by neighboring synapses, a longer axon or the cell body. Photobleached molecules are reinserted as unbleached when moving via the periodic boundary condition. The length of the axon was varied and chosen sufficiently long, so that the axon length and the periodic boundary condition have only a minor effect on the recovery times. Moreover, the chosen length of 5 *µ*m is consistent with experimental data where the mean distance between synapses is similar to the axon length in our simulations [].

### C. FRAP simulations

To simulate FRAP experiments, we first equilibrate our simulations by waiting for the protein concentrations to reach a steady state. Afterwards proteins are “photobleached” according to one of the three protocols and the area in which we monitor the recovery is selected. Photobleaching is represented in our simulation by flagging proteins as bleached if they are in the region of the laser pulse at the time of the pulse (or during the photobleaching time interval in case of non-instantaneous bleaching).

For the protocol that photobleaches the entire synapse, we bleach the synapse instantaneously and over its entire length and also monitor the recovery in this whole region. In the instantaneous localized protocol (“0 ms”), we bleach a 400 nm diameter cylinder in the center of the synapse and use this also as our region of interest (ROI) for monitoring the recovery. In the continuous bleaching (“80 ms”) protocol, we bleach the same 400 nm diameter cylinder in the center of the synapse over a time interval of 80 ms. Throughout this time, every protein that enters the bleaching spot is flagged as photobleached. Subsequently, we choose the region to monitor the recovery by comparing the fluorescence intensity before and after the bleaching, following the protocol for analyzing FRAP experiments from ref. [20]. We create synthetic microscopy images from our simulations by projecting the 3D concentrations of the fluorescent proteins onto a 2D plane and taking a convolution with the experimentally obtained point spread function of GFP to generate a 128x128 pixel image. Similar to the experiments, we generate two images, one immediately before photobleaching and one 0.5 s after photobleaching (i.e., after the start of the laser pulse) and calculate the difference in intensity between two synthetic microscope images *I*_*D*_. From the calculated difference, we select the region of interest (ROI) as all pixels for which the intensity changed significantly. This was determined using the criterion: *I*_*D*_ *>* ⟨*I*_*D*_⟩ + 0.5 std(*I*_*D*_). Impor- tantly, the average and standard deviation of the intensity are calculated over the whole image including the empty/dark background.

Finally, the recovery curves are obtained by summing up the intensity inside the ROI and tracking this sum over time. In the case of the exact protocol we adhered to monitoring at the same frame rate as in the experiments of ref. [20]: first, we measured at intervals of 0.5s for 12 seconds, subsequently at intervals of 1s till 36s after bleaching and finally at intervals of 2s. In this way, we recorded the intensity for 80s. In the other protocols we choose the time resolution to be 0.1s and recorded for 200s.

The resulting curves are fitted to a single exponential *I*(*t*) = *I*_0_(1 − *e*^−*t/*τ^), from which we extract the recovery time τ and also the immobile fraction by comparing the intensity prior to and after bleaching. The exact procedure to obtain FRAP results in the axon is identical except that the criterion to detect the ROIs was *I*_*D*_ *>* ⟨*I*_*D*_⟩ + std(*I*_*D*_).

### D. Fitting FRAP experiments

We first generated a library of simulated FRAP experiments, systematically varying the parameters of the model (*D, k, T*) and determined the recovery times τ_*axon*_, τ_*syn*_ and im- mobile fractions IF_syn_, IF_axon_ for each case. For each protein, we used the corresponding experimental results for recovery times τ_*axon*_, τ_*syn*_ and immobile fractions IF_syn_, IF_axon_ to fit our simulations. First, we regularized the data. To this end, we calculated the mean

*µ*{τ_syn_,τ_*syn*_,*IF*_*syn*_,*IF*_*axon*_} and the standard deviation *σ{*τ_*syn*_,τ_*syn*_,*IF*_*syn*_,*IF*_*axon*_*}* of the distribution of the 14 experimental values. Subsequently, for each protein we subtracted the mean from the original value and divided by the standard deviation e.g. 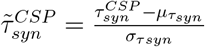. We applied the same transformation to our simulation data. The resulting fit was obtained by subsequently choosing from our library the simulation which had the smallest euclidean distance from the experimental data.

## V. ACKNOWLEDGEMENTS

This work was supported by the Deutsche Forschungsgemeinschaft (DFG) through SFB 1286 (project ID 317475864), projects B02 (to S.R.) and C05 (to S.K.). The simulations were run on the GoeGrid cluster at the University of Göttingen, which is supported by the Deutsche Forschungsgemeinschaft (Project IDs 436382789; 493420525) and MWK Nieder- sachsen (grant no. 45-10-19-F-02).

## SUPPLEMENTARY FIGURES

**FIG 7.**
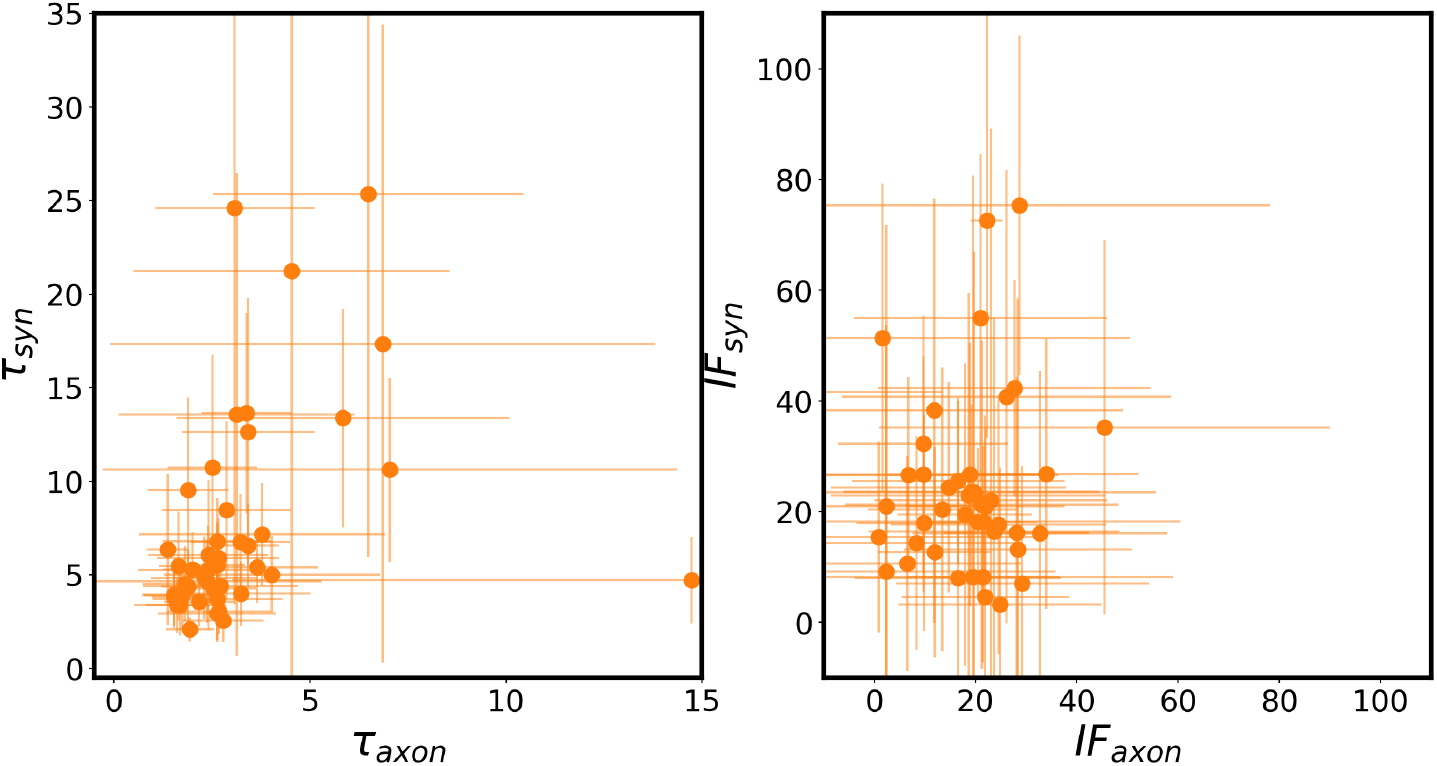
Overview of FRAP results: Recovery times and immobile fractions in the synapse and the axon for 42 proteins (data from ref. [20]).

**FIG 8.**
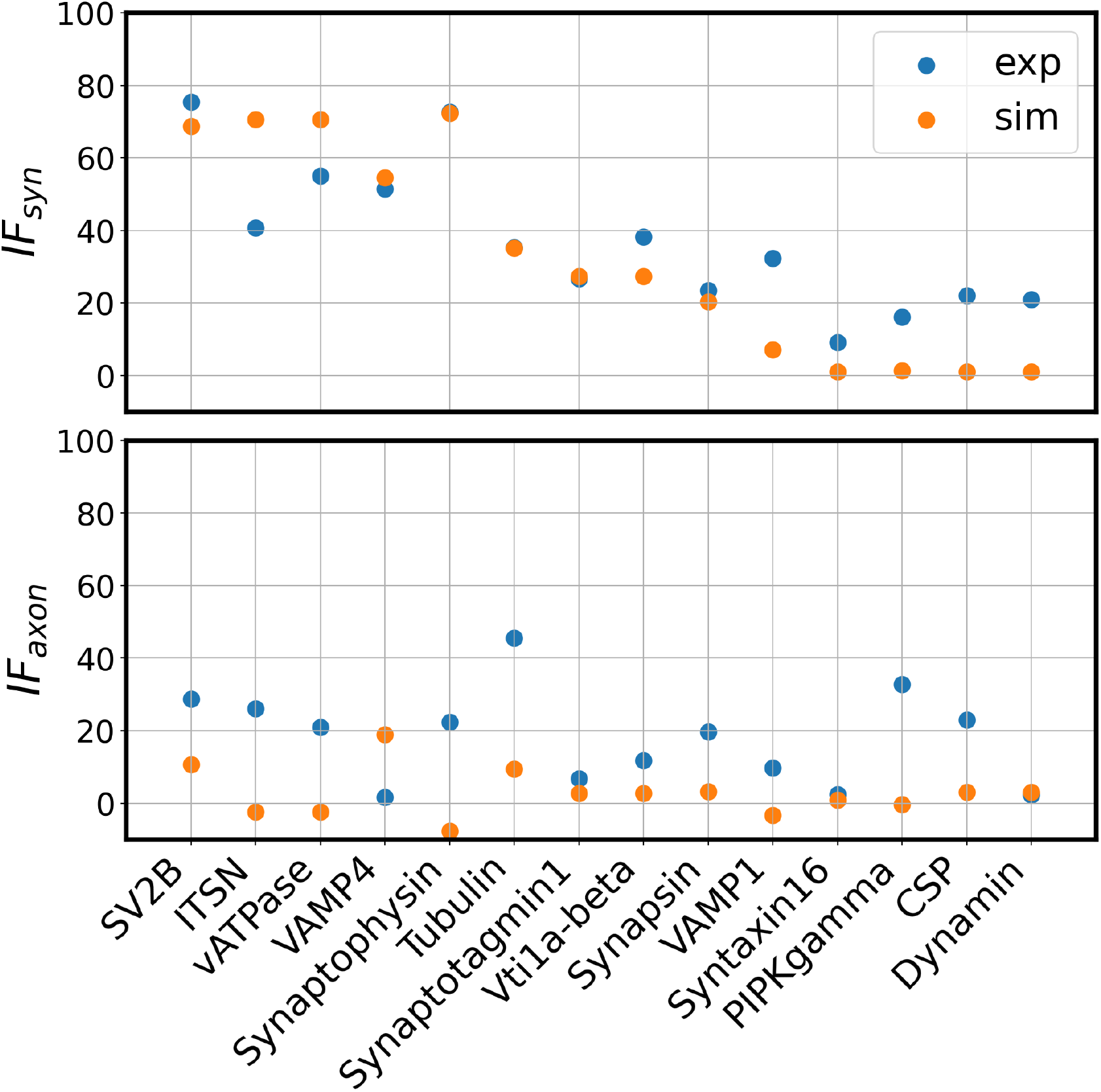
Simulated mmobile fractions (orange) obtained after matching simulations to experimental FRRAP data (blue).

